# Prescribing of different antibiotics, rates of sepsis-related mortality and bacteremia in the US and England, and the utility of antibiotic replacement vs. reduction in prescribing

**DOI:** 10.1101/527101

**Authors:** Edward Goldstein

**Affiliations:** Center for Communicable Disease Dynamics, Department of Epidemiology, Harvard TH Chan School of Public Health, Boston, MA 02115 USA.

## Abstract

**Background:** Antibiotic use contributes to the rates of bacteremia, sepsis and associated mortality, particularly through lack of clearance of resistant infections following antibiotic treatment. At the same time, there is limited information on the effects of prescribing of some antibiotics vs. others on the rates of outcomes related to severe bacterial infections.

**Methods:** We looked at associations (univariate, as well as multivariable for the US data) between the proportions (state-specific in the US, Clinical Commissioning Group (CCG)-specific in England) of different antibiotic types/classes among all prescribed antibiotics in the outpatient setting (oral antibiotics in the US), and rates of outcomes (mortality with sepsis, ICD-10 codes A40-41 present on the death certificate in different age groups of US adults, and *E. coli* and MSSA bacteremia in England) per unit of antibiotic prescribing (defined as the rate of outcome divided by the rate of prescribing of all antibiotics).

**Results:** In the US, prescribing of penicillins was associated with rates of mortality with sepsis for persons aged 75-84y and 85+y between 2014-2015, while multivariable analyses also shown an association between the percent of individuals aged 50-64y lacking health insurance, as well as the percent of individuals aged 65-84y who are African-American and rates of mortality with sepsis. In England, prescribing of penicillins other than amoxicillin/co-amoxiclav was associated with rates of both MSSA and *E. coli* bacteremia for the period between financial years 2014/15 through 2017/18.

**Conclusions:** Our results suggest that prescribing of penicillins is associated with rates of *E. coli* and MSSA bacteremia in England, and rates of mortality with sepsis in older US adults, which agrees with our earlier findings. Those results, as well as the related epidemiological data suggest that replacement of certain antibiotics, particularly penicillins should be considered for reducing the rates of outcomes related to severe bacterial infections.

## Introduction

Rates of hospitalization with septicemia and sepsis in the diagnosis, associated mortality, as well as monetary costs of those hospitalizations have been rising rapidly during the past decades in the US [1–4]. A recent estimate from the US CDC suggests that about 270,000 Americans die annually as a result of sepsis [5]. Moreover, that estimate is expected to increase significantly if longer-term, e.g. 90-day mortality following sepsis diagnosis is accounted for [6]. In England, while rates of certain severe infections and related mortality, such as *Clostridium difficile* infections and MRSA bacteremia have declined during recent years [7,8], rates of *E. coli* and MSSA bacteremia and associated mortality were increasing [7–9].

Bacteremia outcomes in England are laboratory confirmed and there is less uncertainty about the interpretation of the recorded trends for those outcomes compared to trends for septicemia/sepsis hospitalization rates in the US. Part of the reason behind the rapid growth in the rates of hospitalization with septicemia/sepsis in the diagnosis in the US is changes in diagnostic practices, including the implementation of sepsis screening protocols [10,11]. However, changes in diagnostic practices in the US cannot fully explain the rise in the rates of hospitalization with septicemia/sepsis in the diagnosis, particularly prior to 2010 [12]. Indeed, trends in the rates of hospitalizations with any diagnosis of sepsis in the US between 2003-2009 closely resemble the trends in the rates of hospitalizations that involved infection and the use of mechanical ventilation (Figure 1 in [12]). Moreover, rates of hospitalization with severe sepsis in the diagnosis were growing robustly between 2008-2012, with the percent of hospitalizations with severe sepsis that involved multiple organ failure also rising during that period [13], suggesting genuine growth in the volume of hospitalization involving severe sepsis.

Antibiotic use and resistance can contribute to the rates of bacteremia/sepsis hospitalization and mortality through several mechanisms, particularly lack of clearance of resistant infections/colonization following antibiotic treatment, with some of those infections subsequently devolving into bacteremia/sepsis, and lethal outcomes [14–19]. Some of the more direct evidence for the relation between antibiotic resistance and subsequent hospitalization with severe infections, including bacteremia/sepsis is described in [19,18]; evidence about the relation between antibiotic resistance for hospitalized patients with sepsis and mortality, particularly in the US is presented in [16,17]. Those relations suggest that replacement of certain antibiotics by those antibiotics to which prevalence of resistance is lower is expected to help bring down the rates of severe outcomes associated with bacterial infections. For example, prevalence of co-amoxiclav resistance in bacteremia outcomes in England, particularly *E. coli* bacteremia is very high [20] (e.g. more than twice as high as prevalence of co-amoxicalv resistance in *E.* coli-related urinary tract infections [21]), and use of co-amoxiclav [22], both in the hospital and the primary care settings, and possibly the use of related penicillins may contribute to the incidence of co-amoxiclav resistant *E. coli* infections/colonization and associated bacteremia outcomes. We note that guidelines for replacement of certain antibiotics by certain others are relatively less common compared to the recommendation for overall reduction in antibiotic use issued by public health entities in different countries, e.g. [23]. However, reduction in antibiotic prescribing (rather than antibiotic replacement) is less likely to bring down the rates of bacteremia/sepsis in the short term as lack of treatment is generally worse than no treatment in relation to bacteremia/sepsis outcomes. For example, rates of bacteremia kept growing rapidly in England [21] while the rates of antibiotic consumption in the UK dropped by 7.3% from 2014 to 2017 [23]. Moreover, reduction in prescribing, even a relatively modest one, may potentially contribute to increases in the volume of certain outcomes such as pneumonia [24,25]. At the same time, reduction in prescribing may help bring down the rates of severe bacterial infections in the longer term through decrease in antibiotic resistance (e.g. [9]) as antibiotic use is an important driver of the prevalence of antibiotic resistance [26–29,9]. Moreover, antibiotic use may contribute to the prevalence of resistance not only to the drug class used, but to other drugs as well as resistance to different drug classes tends to cluster in bacterial populations, leading to the phenomenon of coselection [30,31]. For example, fluoroquinolone use was found to be associated with methicillin-resistant *S. aureus* (MRSA) infections [32–34], while amoxicillin use was found to be associated with trimethoprim resistance in Enterobacteriaceae in England [26], with trimethoprim resistance in urinary tract infections (UTIs) being positively associated with bacteremia outcomes [27].

There is geographic variability in overall antibiotic prescribing rates within different countries including the US and England, with that variability being associated with variability in the prevalence of underlying health conditions and certain demographic factors [35,36], as well as variability in the rates of severe outcomes associated with bacterial infections [14]-see also Tables 1 and 5 in this paper. However, less is known about the effect of using some antibiotics vs. others in the treatment of various syndromes on the rates of bacteremia, septicemia/sepsis, and associated mortality. Our earlier work [14] studied the relation between the use of different antibiotics and rates of septicemia hospitalization in US adults. In this paper, we examine how the proportions of overall antibiotic prescribing that are for different antibiotic types/classes are related to the rates of *E. coli* and MSSA bacteremia in England, and the rates mortality with sepsis in different age groups of US adults. Those analyses are based on state-level US CDC data on outpatient antibiotic prescribing and mortality between 2014-2015 in [37,38], and on the Clinical Commissioning Groups (CCG)-level English data (from Oxford U/PHE) on GP antibiotic prescribing and bacteremia [39,40]. Additionally, we use a multivariable framework to relate the proportions of overall outpatient antibiotic prescribing that are for fluoroquinolones, penicillins, cephalosporins and macrolides to rates of mortality with sepsis in different age groups of US adults, adjusting for additional covariates and random effects. We hope that such ecological analyses would lead to further work on the effect of antibiotic prescribing, including replacement of some antibiotics by others and reduction in antibiotic prescribing (as well as the comparison between the effect of antibiotic replacement vs. reduction in prescribing – see Discussion) on the rates of bacteremia, sepsis and associated mortality.

## Materials and Methods

### Data

All the data used in this study are publicly available and accessible through refs. [37–40,42–45] as described below.

1. *US.* Data on annual state-specific mortality with sepsis (ICD-10 codes A40-A41.xx representing either the underlying or a contributing cause of death) between 2014-2015 for different age groups of adults (18-49y, 50-64y, 65-74y, 75-84y, 85+y) were extracted from the US CDC Wonder database [38]. For each age group, those data are available for the 50 US states and the District of Columbia (sample size of 51). We note that for most of those deaths, sepsis is listed as a contributing rather than the underlying cause of death on the death certificate [41]. Data on the annual state-specific per capita rates of outpatient prescribing for four classes of oral antibiotics: fluoroquinolones, penicillins, macrolides, and cephalosporins, as well as overall antibiotic prescribing in different states in 2014 and 2015 were obtained from the US CDC Antibiotic Patient Safety Atlas database [37]. Annual state-specific population estimates in each age group of adults (overall, as well as the number of African-Americans) were obtained as the yearly July 1 population estimates in [42]. Data on median household income for US states between 2014-2015 were extracted from [43]. Data on average daily temperature for US states were obtained from [44]. Data on the percent of state residents in different age groups who lacked health insurance were extracted form the US Census Bureau database [45].
2. *England.* We’ve considered the following nine antibiotic types/classes:

1. Amoxicillin (British National Formulary (BNF) code 0501013B0)
2. Co-amoxiclav (Amoxicillin/Clavulanic acid) (BNF code 0501013K0)
3. Penicillins (BNF section 5.1.1) excluding amoxicillin/co-amoxiclav
4. Tetracyclines (BNF section 5.1.3)
5. Macrolides (BNF section 5.1.5)
6. Cephalosporins + other beta-lactams (BNF section 5.1.2)
7. Fluoroquinolones (BNF section 5.1.12)
8. Trimethoprim (BNF code 0501080W0)
9. Urinary Tract Infection antibiotics (BNF 5.1.13 --nitrofurantoin / fosfomycin/methenamine)

For each antibiotic type/class above, we’ve extracted data for the different Clinical Commissioning Groups (CCGs) on the proportion of that antibiotic type/class among all General Practitioner (GP) antibiotic prescriptions (BNF classes 5.1.1 through 5.1.13) in the given CCG for each of the financial years (April through March) 2014/15 through 2017/18 [39]. We’ve also extracted CCG/year specific data on the prescribing of all antibiotics per 1,000 residents, as well as per 1,000 STAR-PUs [39,46]. In addition to prescribing data, we’ve extracted CCG/year-specific data on the (population-adjusted) rates of *E. coli* and MSSA bacteremia for each of the financial years 2014/15 through 2017/18 [40]. We note that mergers of certain CCGs took place during the study period, and not all CCGs reported all the data needed for our analyses. For the 197 CCGs that reported data in [41], we have included data on 189 CCGs that reported both annual data on *E. coli* and MSSA bacteremia, as well as data on prescribing of the nine antibiotic types/classes above for each of the financial years 2014/15 through 2017/18.

### Univariate Correlations (US and England)

The contribution of prescribing of a given antibiotic type/class to the rates of severe outcomes associated with bacterial infections (e.g. bacteremia, or mortality with sepsis) is expected to be proportional to the rate of prescribing of that antibiotic type/class. One of the factors that modulates the relationship between the rates of antibiotic prescribing and rate of severe outcomes is the rate of infection that affects both prescribing and severe outcomes. Correspondingly, associations between rates of antibiotic prescribing and rates of severe outcomes associated with bacterial infections are often positive (e.g. Tables 1 and 5 in this paper). Moreover, the use of certain antibiotics may have a stronger association with severe outcomes than the use of certain other antibiotics, e.g. due to differences in the prevalence of resistance to different antibiotics. If a unit of prescribing of a given antibiotic type/class has a stronger association with the rate of a given severe outcome compared to prescribing of an average antibiotic dose (e.g. as a result of high prevalence of resistance to a given antibiotic), the association between the *proportion* of given antibiotic type/class among all antibiotics prescribed and the rate of a given severe outcome *per unit of antibiotic prescribing* (defined as the rate of severe outcomes divided by the rate of prescribing of all antibiotics) is expected to be positive. We note that such disproportionate effects of prescribing of a unit of a given antibiotic can be the result not only of treatment of infections leading to a given outcome by a given antibiotic, but also of the contribution of the use of a given antibiotic to the rates of infection/colonization with different bacteria, and the prevalence of resistance to other antimicrobials (see Introduction). Additionally, correlations between proportions of a given antibiotic type/class among all oral antibiotics prescribed and the rates of severe outcomes per unit of antibiotic prescribing can also be affected by patterns of antibiotic prescribing in different locations, including changes in those patterns resulting from increases in resistance, introduction of new prescribing guidelines, etc. Correspondingly, we studied correlations between the proportions (state-specific in the US, CCG-specific in England) of a given antibiotic type/class among all prescribed antibiotics in the outpatient setting, and rates of outcome (mortality with sepsis in different age groups of adults in the US, and *E. coli,* as well as MSSA bacteremia in England) per unit of antibiotic prescribing, with the caveats above regarding the causal relations underlying those correlations. The US analysis is done for the 2014-2015 period; given the ongoing changes in antibiotic prescribing patterns in England [23], correlations for the English data were computed for each financial year between 2014/15 through 2017/18.

### Multivariable model (US data)

In this section, we apply a mixed-effect multivariable model to adjust for various factors that affect the relation between prescribing of different antibiotics and rates of severe outcomes (including mortality) associated with bacterial infections. The relevant data for a multivariable model were only available for the US. For each age group of adults, (18-49y, 50-64y, 65-74y, 75-84y, 85+y), we applied mixed effect models to relate the average annual state-specific outpatient prescribing rates (per 1,000 state residents) for oral fluoroquinolones, penicillins, macrolides, and cephalosporins between 2014-2015 to the average annual state-specific rates of sepsis mortality per 100,000 individuals in a given age group between 2014-2015 (dependent variable). Besides the antibiotic prescribing rates, the other covariates were the state-specific median household income, percentages of state residents in a given age group who were African American, those who lacked health insurance (in the non-elderly age groups, as health insurance, particularly Medicare coverage levels in the elderly are very high), as well as the state-specific average annual temperature. We note that sepsis mortality rates in African Americans are elevated [47]. We also note that temperature may influence bacterial growth rates and/or transmission mediated effects [48], which in turn may affect both the prevalence of antibiotic resistance [48], and the acquisition/severity of bacterial infections. To adjust for additional factors not accounted for by the covariates used in the model, we include random effects for the ten Health and Human Services (HHS) regions in the US. Specifically, for each state *s,* let *MR* (*s*’) be the average annual state-specific rate of mortality (per 100,000) with sepsis in the given age group between 2014-2015, *A_i_*(*s*) (*i* = 1,..,4) be the average annual state-specific outpatient prescribing rates, per 1,000 state residents (of all ages), for the four studied classes of antibiotics between 2014-2015 (thus *A*_1_(*s*) denotes the rate of prescribing of oral fluoroquinolones, etc.); *I*(*s*) be the median state-specific household income between 2014-2015; T(s) be the state-specific average annual temperature (°F) between 2002-2011; *AA*(*s*) be the age-specific percent of state residents between 2014-2015 who were African American; *LHI*(*s*) be the average annual age-specific percent of state residents who lacked health insurance between 2014-2015 (for non-elderly age groups); *α*(*s*) be the random effect for the corresponding HHS region, and *ε* be the residual. Then

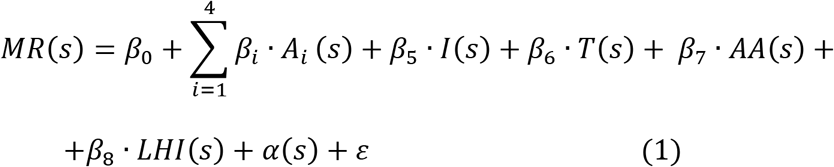

We note that is we divide eq. 1 by the state-specific rates of overall outpatient prescribing of oral antibiotics, the resulting equation expresses (models) a linear relation between proportions of the overall antibiotic prescribing that are for fluoroquinolones, penicillins, macrolides, and cephalosporins, and other covariates and sepsis mortality rates per unit of antibiotic prescribing. Thus, eq. 1 can be thought of as a multivariable model for the relation between proportions of overall oral antibiotic prescribing that are for given antibiotic types/classes and rates of outcomes associated with bacterial infections per unit of oral antibiotic prescribing, with a univariate model for those relations studied in the previous subsection of the Methods.

## Results

### 1. US

Table 1 shows, for each age group of adults, the mean (standard deviation) for the state-specific average annual rates of mortality with sepsis per 100,000 individuals in that age group between 2014-2015, as well as the linear correlation between those rates and state-specific rates of outpatient prescribing of all oral antibiotics. The latter correlations are high, ranging from 0.59(0.37,0.74) for ages 85+y to 0.77(0.62,0.68) for ages 65-74y. Additionally, annual rates of mortality with sepsis increase rapidly with age, from the state-specific mean of 8.31/100,000 for persons aged 18-49y to a mean of 750/100,000 for persons aged 85+y.

**Table 1:** State-specific rates of mortality with sepsis (ICD-10 codes A40-41.xx present as either underlying or contributing causes on a death certificate) per 100,000 individuals in different age groups between 2014-2015 (mean + standard deviation), and the linear correlation between those rates and state-specific rates of outpatient prescribing of all oral antibiotics.

|  | Mean<br>(standard deviation) | Linear correlation with rate of<br>prescribing of all oral antibiotics |
| --- | --- | --- |
| Sepsis mortality rate,<br>ages 18-49y | 8.31 (2.98) | **0.66(0.47,0.79)** |
| Sepsis mortality rate,<br>ages 50-64y | 55.8 (17.3) | **0.74(0.59,0.84)** |
| Sepsis mortality rate,<br>ages 65-74y | 143.4 (36.5) | **0.77(0.62,0.68)** |
| Sepsis mortality rate,<br>ages 75-84y | 330.8 (75.6) | **0.67(0.48,0.80)** |
| Sepsis mortality rate,<br>ages 85+y | 750 (161.9) | **0.59(0.37,0.74)** |

Table 2 shows correlations (both linear and Spearman) for each pair of antibiotic classes between the state-specific percentages of all outpatient oral antibiotic prescriptions that were for each antibiotic class between 2014-2015, as well as the mean (standard deviation) for the state-specific percentages of all outpatient oral antibiotic prescriptions that were for each antibiotic class between 2014-2015. There is a strong negative correlation between the percentages of outpatient prescribing of oral antibiotics that are for fluoroquinolones and that are for penicillins; the Spearman correlation between percentages of antibiotic prescribing that are for penicillins and that are for cephalosporins is also negative. Those negative correlations suggest competition between certain antibiotics in outpatient prescribing for various syndromes. We also note that on average, 66.2% of all outpatient oral antibiotic prescriptions in different states were for the four studied classes of antibiotics. Additionally, proportions of different antibiotic classes among all oral antibiotics prescribed in the outpatient setting in the US are notably different from the corresponding proportions in England (compare Table 2 with Table 6). Those differences, particularly for fluoroquinolones and cephalosporins may be related to differences in the rates of severe bacterial infections, particularly *Clostridium difficile* (C. difficile) infections between the two countries [49], with reduction in fluoroquinolone and cephalosporin prescribing found to be associated with reduction in the incidence of C. difficile infection in both countries [50,51].

**Table 2:** Correlations (both linear (Pearson, with 95% confidence intervals) and Spearman (with p-values)) between state-specific proportions of prescriptions for each antibiotic class among all outpatient oral antibiotic prescriptions in the state for different pairs of antibiotic classes, as well as the mean (standard deviation) for the average annual state-specific percentages of all antibiotic prescriptions that were for each antibiotic class between 2014-2015.

|  | Mean<br>(standard<br>deviation) | Linear (Pearson) correlation with<br>proportions of other antibiotic classes |  |  |
| --- | --- | --- | --- | --- |
|  |  | Proportion<br>penicillins | Proportion<br>cephalosporins | Proportion<br>Macrolides |
| Proportion<br>fluoroquinolones | 11.81%<br>(1.22%) | **-0.563**<br>**(-0.73,-0.34)** | -0.141<br>(-0.4,0.14) | 0.123<br>(-0.16,0.39) |
| Proportion<br>Penicillins | 22.57%<br>(1.64%) |  | -0.262<br>(-0.5,0.01) | -0.235<br>(-0.48,0.04) |
| Proportion<br>cephalosporins | 13.56%<br>(1.75%) |  |  | -0.164<br>(-0.42,0.12) |
| Proportion<br>macrolides | 18.22%<br>(1.21%) |  |  |  |

**Table 2:** Correlations (both linear (Pearson, with 95% confidence intervals) and Spearman (with p-values)) between state-specific proportions of prescriptions for each antibiotic class among all outpatient oral antibiotic prescriptions in the state for different pairs of antibiotic classes, as ...
|  | Mean<br>(standard<br>deviation) | Spearman correlation with proportions of<br>other antibiotic classes |  |  |
| --- | --- | --- | --- | --- |
|  |  | Proportion<br>penicillins | Proportion<br>cephalosporins | Proportion<br>Macrolides |
| Proportion<br>fluoroquinolones | 11.81%<br>(1.22%) | **-0.436**<br>**(0.0015)** | -0.069<br>(0.63) | 0.114<br>(0.42) |
| Proportion<br>penicillins | 22.57%<br>(1.64%) |  | **-0.306**<br>**(0.03)** | -0.192<br>(0.18) |
| Proportion<br>cephalosporins | 13.56%<br>(1.75%) |  |  | -0.197<br>(0.16) |

Table 3 shows correlations (both Spearman and linear), for each antibiotic class and age group, between average annual state-specific proportions of a given antibiotic class among the overall outpatient oral antibiotic prescriptions and average annual state-specific rates of mortality with sepsis in a given age group per unit of prescribed antibiotics (Methods) between 2014-2015. The Spearman correlations are positive for penicillins for persons aged 75-84y and over 85y, and for fluoroquinolones for persons aged 50-64y; the Spearman correlations are negative for cephalosporins for persons aged 75-84y and over 85y, and for penicillins for persons aged 18-49y. Among those six significant Spearman correlations, all the corresponding linear correlations are also significant save for penicillins and mortality with sepsis in persons aged 75-84y (see also Table 4).

**Table 3:** Correlations (both Spearman (p-value) and linear (Pearson, 95% CI)) between average annual state-specific percent of overall outpatient oral antibiotic prescribing that is for a given antibiotic class and average annual state-specific rates of mortality with sepsis in a given age group per unit of prescribed oral antibiotics (Methods) between 2014-2015.

|  |  | Proportion<br>fluoroquinolones | Proportion<br>penicillins | Proportion<br>cephalosporins | Proportion<br>macrolides |
| --- | --- | --- | --- | --- | --- |
| Age<br>18-49y | Spearman | 0.219<br>(0.12) | **-0.36**<br>**(0.01)** | 0.262<br>(0.064) | 0.235<br>(0.097) |
|  | Linear | 0.142<br>(-0.14,0.4) | **-0.349**<br>**(-0.57,-0.08)** | 0.168<br>(-0.11,0.42) | 0.269<br>(-0.01,0.51) |
| Age<br>50-64y | Spearman | **0.419**<br>**(0.002)** | -0.233<br>(0.1) | -0.063<br>(0.66) | 0.185<br>(0.19) |
|  | Linear | **0.35**<br>**(0.08,0.57)** | -0.183<br>(-0.44,0.1) | -0.151<br>(-0.41,0.13) | 0.205<br>(-0.07,0.46) |
| Age<br>65-74y | Spearman | 0.274<br>(0.052) | 0.003<br>(0.98) | -0.247<br>(0.08) | 0.092<br>(0.52) |
|  | Linear | 0.258<br>(-0.02,0.5) | -0.011<br>(-0.29,0.27) | **-0.306**<br>**(-0.54,-0.03)** | 0.109<br>(-0.17,0.37) |
| Age<br>75-84y | Spearman | 0.081<br>(0.57) | **0.292**<br>**(0.038)** | **-0.418**<br>**(0.002)** | -0.004<br>(0.98) |
|  | Linear | 0.028<br>(-0.25,0.3) | 0.229<br>(-0.05,0.47) | **-0.411**<br>**(-0.62,-0.15)** | 0.028<br>(-0.25,0.3) |
| Age<br>85+y | Spearman | -0.023<br>(0.87) | **0.416**<br>**(0.003)** | **-0.422**<br>**(0.002)** | -0.048<br>(0.74) |
|  | Linear | -0.04<br>(-0.31,0.24) | **0.354**<br>**(0.09,0.57)** | **-0.442**<br>**(-0.64,-0.19)** | -0.024<br>(-0.3,0.25) |

Table 4 shows the results of the multivariable model given by eq. 1. Table 4 suggests positive associations (with largest effect size in the corresponding age groups) between rates of outpatient prescribing of oral penicillins and sepsis mortality rates in individuals aged 75-84y and 85+y, and a negative association between rates of outpatient prescribing of oral penicillins and sepsis mortality rates in individuals aged 18-49y, all of which agree with the univariate results in Table 3. Table 4 also suggests a positive association between the rates of outpatient prescribing of oral cephalosporins and sepsis mortality rates in individuals aged 18-49y (with the corresponding association failing to reach statistical significance in the univariate model), as well as positive associations between the percent of individuals aged 50-64y lacking health insurance, as well as the percent of individuals aged 65-74y and 75-84y who were African-American and rates of mortality with sepsis.

**Table 4:** Regression coefficients for the different covariates in the model given by eq. 1 for different age groups. The coefficients for the different antibiotic classes estimate the change in the annual sepsis mortality rates (per 10,000 individuals in a given age group) when the annual rate of outpatient prescribing of oral antibiotics in the corresponding class (per 1,000 residents) increases by 1. ND=not done because persons aged >64 years old are eligible for Medicare.

|  | Aged<br>18-49y | Aged<br>50-64y | Aged<br>65-74y | Aged<br>75-84y | Aged<br>85+y |
| --- | --- | --- | --- | --- | --- |
| Fluoroquinolones<br>(prescription per<br>1000 residents/y) | 0.01<br>(-0.04,0.07) | 0.15<br>(-0.15,0.45) | 0.26<br>(-0.4,0.92) | -0.16<br>(-1.77,1.45) | -0.61<br>(-4.52,3.3) |
| Penicillins<br>(prescription per<br>1000 residents/y) | **-0.03**<br>**(-0.07,0)** | 0.08<br>(-0.1,0.25) | 0.11<br>(-0.28,0.5) | **0.95**<br>**(0.02,1.88)** | **2.97**<br>**(0.72,5.22)** |
| Cephalosporins<br>(prescription per<br>1000 residents/y) | **0.05**<br>**(0.02,0.09)** | 0.07<br>(-0.11,0.25) | 0.13<br>(-0.28,0.55) | -0.06<br>(-1.04,0.93) | -0.76<br>(-3.09,1.58) |
| Macrolides<br>(prescription per<br>1000 residents/y) | 0.02<br>(-0.02,0.06) | 0.06<br>(-0.15,0.26) | 0.21<br>(-0.26,0.69) | 0.45<br>(-0.69,1.58) | 0.71<br>(-2.03,3.45) |
| Median household<br>income (\$1000) | -0.06<br>(-0.13,0.01) | -0.17<br>(-0.55,0.2) | -0.09<br>(-0.9,0.73) | 0.54<br>(-1.4,2.49) | 1.95<br>(-2.7,6.59) |
| Average minimal daily<br>temperature (°F) | -0.04<br>(-0.11,0.03) | 0.16<br>(-0.19,0.51) | 0.33<br>(-0.44,1.1) | 0.6<br>(-1.26,2.46) | 4.07<br>(-0.55,8.7) |
| Percent African<br>Americans | 0.03<br>(-0.05,0.11) | 0.28<br>(-0.06,0.63) | **1.25**<br>**(0.41,2.1)** | **2.68**<br>**(0.76,4.6)** | 3.02<br>(-1.63,7.66) |
| Percent lacking health<br>insurance | 0.05<br>(-0.1,0.2) | **0.95**<br>**(0.01,1.89)** | ND | ND | ND |

### 2. England

Table 5 shows summary statistics (mean + standard error) for the annual rates of *E. coli* and MSSA bacteremia for the different CCGs for the 2014/15 through the 2017/18 financial years, as well as the correlation between those rates and the rates of GP antibiotic prescribing, both per 1,000 residents and per 1,000 STAR-PUs [46]. Table 5 suggests an ongoing increase in the rates of MSSA bacteremia [7], with the long-term growth in the rates of *E. coli* bacteremia [9,7,21,40] stalling in 2017/18. Table also 5 shows that for each of the 4 years in the data, for both the *E. coli* and MSSA bacteremia rates, estimates of the correlation between those rates and antibiotic prescribing per 1,000 residents are higher than the estimates of the correlation between those rates and antibiotic prescribing per 1,000 STAR-Pus, suggesting that rates of severe bacterial infections are reflected better by the actuality of antibiotic prescribing in England rather than by the recommendations set by the STAR-PU system.

**Table 5:**
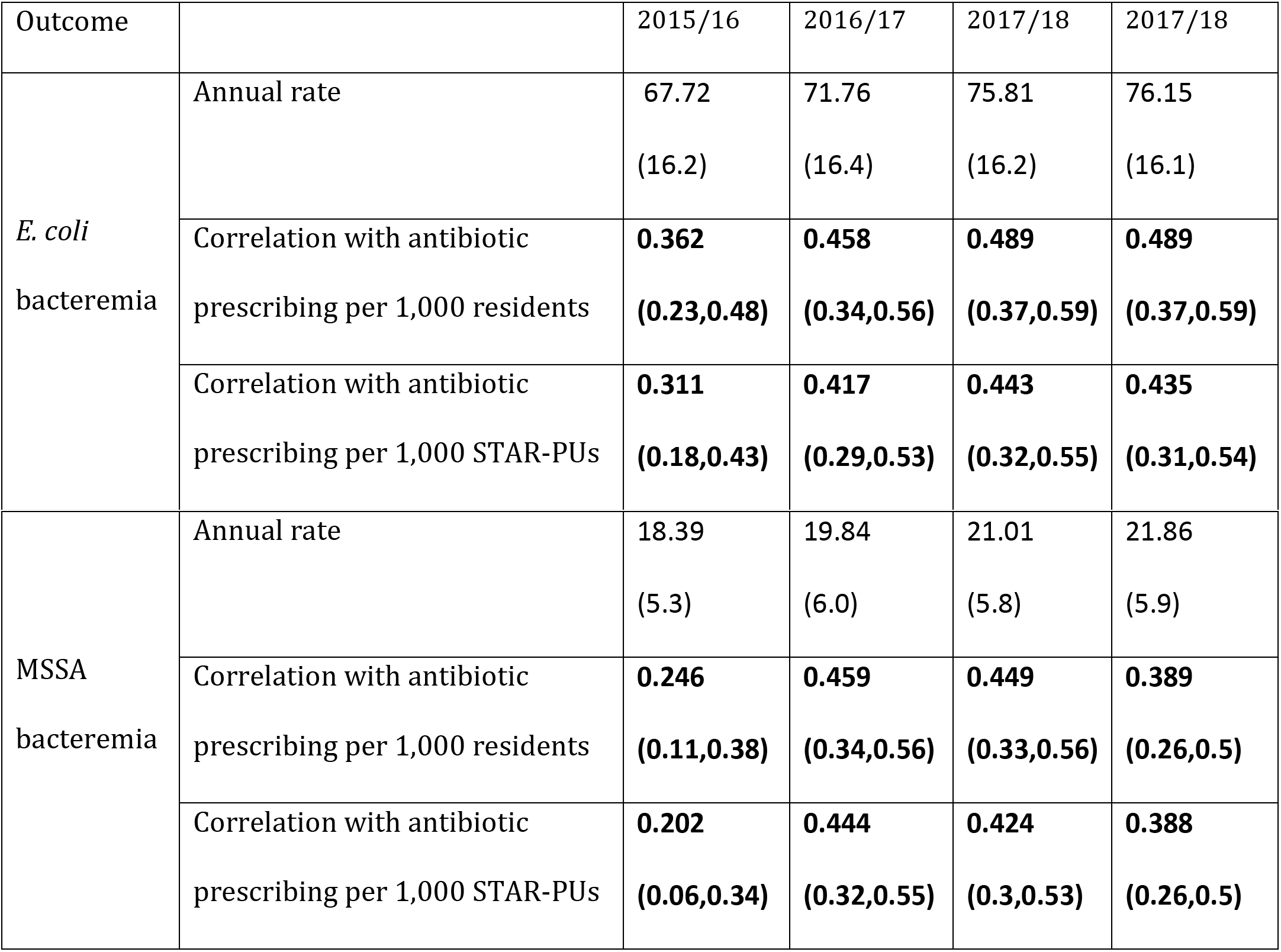
Annual rates of *E. coli* and MSSA bacteremia for the different CCGs (mean + standard error) for the 2014/15 through the 2017/18 financial years; correlations between those bacteremia rates and rates of overall GP antibiotic prescribing, both per 1,000 residents and per 1,000 STAR-PUs [46].

Table 6 shows the mean + standard error for the annual CCG-specific rates of GP prescribing of different antibiotic types/classes per 1,000 residents, as well as for the annual CCG-specific proportions of those antibiotic types/classes among all prescribed antibiotics for the 2014/15 through the 2017/18 financial years. Table 6 suggests substantial temporal reduction in the rates/proportions of prescribing for trimethoprim, co-amoxiclav and cephalosporins/other beta lactams, as well as reduction in the rates/proportions of amoxicillin, fluroquinolone and macrolide prescribing. Prescribing of UTI antibiotics (BNF 5.1.13 -- nitrofurantoin/fosfomycin/methenamine) increased markedly (with a good amount of replacement of trimethoprim by nitrofurantoin in the treatment of UTIs taking place during the study period, [20]), with proportions of penicillins other than amoxicillin/co-amoxiclav and tetracyclines among all prescribed antibiotics also increasing. Additionally, significant reduction in trimethoprim prescribing took place in 2017/2018 compared to 2016/17, and the growth in the rate of *E. coli* bacteremia had also stalled then (Tables 6 and 5).

**Table 6:** Annual CCG-specific proportions (percentages) of a given antibiotic type/class among all GP antibiotic prescriptions (mean + standard error), and annual CCG-specific rates of GP prescribing per 1,000 individuals (mean + standard error) for nine antibiotic types/classes (Methods) during the 2014/15 through the 2017/18 financial years in England.

|  | 2014/15 |  | 2015/16 |  | 2016/17 |  | 2017/18 |  |
| --- | --- | --- | --- | --- | --- | --- | --- | --- |
|  | Percent<br>of all abx | Rate<br>per<br>1,000 | Percent<br>of all abx | Rate<br>per<br>1,000 | Percent<br>of all abx | Rate<br>per<br>1,000 | Percent<br>of all abx | Rate<br>per<br>1,000 |
| Amoxicillin | 28.35%<br>(3.8%) | 187.6<br>(36.3) | 26.8%<br>(3.7%) | 162.9<br>(32.5) | 26.75%<br>(3.5%) | 161.1<br>(31.4) | 26.04%<br>(3.4%) | 150<br>(29.7) |
| Co-amoxiclav | 5.44%<br>(2.2%) | 35.4<br>(14.6) | 4.8%<br>(1.9%) | 28.8<br>(11.4) | 4.28%<br>(1.5%) | 25.4<br>(9.4) | 4.16%<br>(1.4%) | 23.6<br>(8.4) |
| Penicillins except<br>amoxicillin/co-amoxiclav | 17.67%<br>(1.5%) | 116.7<br>(18.1) | 18.41%<br>(1.6%) | 111.8<br>(17.9) | 18.6%<br>(1.7%) | 111.8<br>(18.4) | 19.18%<br>(1.8%) | 110.4<br>(18.9) |
| Tetracyclines | 11.59%<br>(2.9%) | 77.5<br>(23.7) | 12.24%<br>(2.9%) | 75.4<br>(23) | 12.74%<br>(3%) | 77.9<br>(24.1) | 13.12%<br>(3%) | 77<br>(24) |
| Macrolides | 12.66%<br>(1.6%) | 83.9<br>(16.6) | 12.3%<br>(1.5%) | 75.1<br>(15.3) | 12.19%<br>(1.5%) | 73.7<br>(15.4) | 11.94%<br>(1.4%) | 69.1<br>(14.8) |
| Cephalosporins + other<br>beta-lactams | 3.3%<br>(1.4%) | 21.9<br>(10.3) | 2.96%<br>(1.2%) | 18.1<br>(8.1) | 2.7%<br>(1.1%) | 16.4<br>(7.5) | 2.6%<br>(1.1%) | 15.2<br>(7.2) |
| Fluoroquinolones | 1.96%<br>(0.6%) | 12.8<br>(3.8) | 1.9%<br>(0.5%) | 11.4<br>(3.3) | 1.87%<br>(0.5%) | 11.1<br>(3.1) | 1.89%<br>(0.5%) | 10.8<br>(3) |
| Trimethoprim | 10.33%<br>(1.7%) | 68.9<br>(16.6) | 10.48%<br>(1.9%) | 64.5<br>(17) | 9.8%<br>(2.1%) | 59.9<br>(17.7) | 7.3%<br>(1.8%) | 42.8<br>(14.3) |
| UTI antibiotics | 5.77%<br>(1.4%) | 38<br>(9.7) | 7.01%<br>(1.7%) | 42.3<br>(10.3) | 8.05%<br>(2%) | 48.1<br>(12.1) | 10.67%<br>(2.1%) | 61.1<br>(13.9) |

Tables 7 and 8 show correlations (both linear and Spearman) between CCG-specific proportions of different antibiotic types/classes among all GP antibiotic prescriptions and rates of *E. coli* (Table 7) and MSSA (Table 8) bacteremia per unit of antibiotic prescribing (Methods) for the 2014/15 through 2017/18 financial years. For penicillins other than amoxicillin/co-amoxiclav, correlations with rates of MSSA bacteremia were positive for all years, and correlations with rates of *E. coli* bacteremia were positive for the 2014/15 through 2016/17 financial years. For macrolides and fluoroquinolones, the corresponding correlations were generally negative. Correlations with bacteremia rates increased with time for proportions of UTI antibiotics, cephalosporins, and, to a smaller extent, amoxicillin prescribing; the corresponding correlations declined for proportions of trimethoprim and co-amoxiclav prescribing, with all those relative changes presumably related more to changes in prescribing patterns rather than changes in the causal relation between the use of a unit of those antibiotics and bacteremia outcomes. In particular, Tables 7 and 8 suggest that relative reductions in trimethoprim and co-amoxiclav prescribing were greater in places with higher bacteremia rates compared to places with lower bacteremia rates. Finally, we note that positive correlations with bacteremia rates for proportions of prescribing for penicillins other than amoxicillin/co-amoxiclav, but not for proportions of co-amoxiclav or amoxicillin prescribing need not suggest that penicillins other than amoxicillin/co-amoxiclav have a stronger relative impact on bacteremia rates than co-amoxiclav or amoxicillin; those differences may also have to do with geographic/demographic variation in the choice of different antibiotics, particularly penicillins.

**Table 7:** Correlations (both linear, with 95% CI, and Spearman, with p-value) between annual proportions of different antibiotic types/classes among all GP antibiotic prescriptions and annual rates of *E. coli* bacteremia per unit of antibiotic prescribing (Methods) for the different CCGs in England, 2014/15 through 2017/18 financial years.

|  | 2014/15 |  | 2015/16 |  | 2016/17 |  | 2017/18 |  |
| --- | --- | --- | --- | --- | --- | --- | --- | --- |
|  | Linear | Spearman | Linear | Spearman | Linear | Spearman | Linear | Spearman |
| Amoxicillin | -0.058<br>(-0.2,0.09) | -0.048<br>(0.5) | 0.017<br>(-0.13,0.16) | 0.036<br>(0.62) | 0.01<br>(-0.13,0.16) | 0.017<br>(0.82) | 0.02<br>(-0.12,0.16) | -0.007<br>(0.93) |
| Co-amoxiclav | -0.056<br>(-0.2,0.09) | -0.056<br>(0.44) | -0.118<br>(-0.26,0.03) | **-0.146**<br>**(0.044)** | -0.14<br>(-0.28,0) | **-0.169**<br>**(0.02)** | -0.14<br>(-0.28,0) | **-0.175**<br>**(0.016)** |
| Penicillins except amoxicillin/<br>co-amoxiclav | **0.256**<br>**(0.12,0.38)** | **0.201**<br>**(0.006)** | **0.204**<br>**(0.06,0.34)** | **0.167**<br>**(0.02)** | **0.15**<br>**(0,0.28)** | 0.117<br>(0.11) | 0.09<br>(-0.05,0.23) | 0.06<br>(0.41) |
| Tetracyclines | **0.171**<br>**(0.03,0.31)** | **0.201**<br>**(0.006)** | 0.095<br>(-0.05,0.23) | 0.115<br>(0.12) | 0.09<br>(-0.06,0.23) | 0.104<br>(0.15) | -0.03<br>(-0.17,0.11) | 0.02<br>(0.78) |
| Macrolides | **-0.208**<br>**(-0.34,-0.07)** | **-0.248**<br>**(0.0006)** | -0.14<br>(-0.28,0) | **-0.156**<br>**(0.03)** | -0.13<br>(-0.27,0.01) | **-0.153**<br>**(0.035)** | -0.07<br>(-0.21,0.08) | -0.091<br>(0.22) |
| Cephalosporins +<br>other beta-lactams | -0.029<br>(-0.17,0.11) | 0.023<br>(0.75) | 0.03<br>(-0.11,0.17) | 0.115<br>(0.11) | 0.02<br>(-0.12,0.16) | 0.039<br>(0.59) | 0.12<br>(-0.02,0.26) | **0.15**<br>**(0.039)** |
| Fluoroquinolones | **-0.276**<br>**(-0.4,-0.14)** | **-0.306**<br>**(0.00002)** | **-0.207**<br>**(-0.34,-0.07)** | **-0.242**<br>**(0.0008)** | **-0.18**<br>**(-0.32,-0.04)** | **-0.203**<br>**(0.005)** | -0.12<br>(-0.26,0.02) | **-0.162**<br>**(0.026)** |
| Trimethoprim | -0.089<br>(-0.23,0.05) | -0.076<br>(0.30) | **-0.279**<br>**(-0.41,-0.14)** | **-0.26**<br>**(0.0003)** | **-0.28**<br>**(-0.41,-0.15)** | **-0.235**<br>**(0.001)** | **-0.26**<br>**(-0.39,-0.12)** | **-0.273**<br>**(0.0002)** |
| UTI antibiotics | 0.038<br>(-0.1,0.18) | -0.065<br>(0.38) | **0.153**<br>**(0.01,0.29)** | 0.044<br>(0.55) | **0.21**<br>**(0.07,0.34)** | 0.135<br>(0.063) | **0.19**<br>**(0.05,0.33)** | **0.164**<br>**(0.024)** |

**Table 8:** Correlations (both linear, with 95% CI, and Spearman, with p-value) between annual proportions of different antibiotic types/classes among all GP antibiotic prescriptions and annual rates of MSSA bacteremia per unit of antibiotic prescribing (Methods) for different CCGs in England, 2014/15 through 2017/18 financial years.

|  | 2014/15 |  | 2015/16 |  | 2016/17 |  | 2017/18 |  |
| --- | --- | --- | --- | --- | --- | --- | --- | --- |
|  | Linear | Spearman | Linear | Spearman | Linear | Spearman | Linear | Spearman |
| Amoxicillin | -0.071<br>(-0.21,0.07) | -0.121<br>(0.10) | -0.047<br>(-0.19,0.1) | -0.076<br>(0.30) | -0.04<br>(-0.18,0.1) | -0.081<br>(0.27) | 0.03<br>(-0.11,0.17) | -0.009<br>(0.90) |
| Co-amoxiclav | -0.114<br>(-0.25,0.03) | -0.112<br>(0.13) | **-0.177**<br>**(-0.31,-0.04)** | **-0.193**<br>**(0.008)** | **-0.26**<br>**(-0.39,-0.12)** | **-0.313**<br>**(0.00001)** | **-0.29**<br>**(-0.42,-0.16)** | **-0.282**<br>**(0.00009)** |
| Penicillins except amoxicillin/<br>co-amoxiclav | **0.259**<br>**(0.12,0.39)** | **0.261**<br>**(0.0003)** | **0.218**<br>**(0.08,0.35)** | **0.172**<br>**(0.018)** | **0.2**<br>**(0.06,0.33)** | **0.196**<br>**(0.007)** | **0.26**<br>**(0.12,0.39)** | **0.232**<br>**(0.001)** |
| Tetracyclines | 0.135<br>(-0.01,0.27) | **0.157**<br>**(0.032)** | **0.223**<br>**(0.08,0.35)** | **0.253**<br>**(0.0005)** | 0.14<br>(0,0.28) | **0.234**<br>**(0.001)** | 0<br>(-0.14,0.15) | 0.057<br>(0.43) |
| Macrolides | **-0.155**<br>**(-0.29,-0.01)** | **-0.183**<br>**(0.012)** | **-0.183**<br>**(-0.32,-0.04)** | **-0.217**<br>**(0.003)** | **-0.17**<br>**(-0.31,-0.03)** | **-0.206**<br>**(0.005)** | -0.13<br>(-0.27,0.01) | **-0.151**<br>**(0.038)** |
| Cephalosporins +<br>other beta-lactams | -0.044<br>(-0.19,0.1) | -0.01<br>(0.89) | -0.046<br>(-0.19,0.1) | -0.024<br>(0.74) | 0<br>(-0.14,0.15) | 0.028<br>(0.71) | 0.04<br>(-0.1,0.18) | 0.075<br>(0.31) |
| Fluoroquinolones | -0.1<br>(-0.24,0.04) | -0.084<br>(0.25) | **-0.206**<br>**(-0.34,-0.07)** | **-0.232**<br>**(0.001)** | -0.12<br>(-0.26,0.02) | **-0.18**<br>**(0.01)** | -0.12<br>(-0.26,0.03) | **-0.148**<br>**(0.04)** |
| Trimethoprim | 0.005<br>(-0.14,0.15) | 0.017<br>(0.82) | **-0.166**<br>**(-0.3,-0.02)** | -0.132<br>(0.07) | **-0.16**<br>**(-0.3,-0.02)** | -0.101<br>(0.17) | **-0.21**<br>**(-0.34,-0.07)** | **-0.214**<br>**(0.003)** |
| UTI antibiotics | 0.001<br>(-0.14,0.14) | -0.029<br>(0.69) | 0.09<br>(-0.05,0.23) | 0.022<br>(0.77) | **0.16**<br>**(0.02,0.3)** | 0.069<br>(0.35) | 0.12<br>(-0.02,0.26) | 0.111<br>(0.13) |

## Discussion

Rates of mortality related to septicemia/sepsis in the US, as well as rates of *E. coli* bacteremia and associated mortality in England are high [1–3,5,7,8,21,40], and antimicrobial use may affect those rates through a variety of mechanisms (Introduction). At the same time, our understanding of the effect of the use of certain antibiotics vs. others for various indications on the rates of bacteremia, septicemia/sepsis and associated mortality is still limited. Additionally, use of certain antibiotics may affect the rates of severe outcomes associated with syndromes for which a given antibiotic is rarely prescribed as use of antibiotics may affect prevalence of infection/colonization and resistance to different antibiotics in different bacterial pathogens that subsequently cause various syndromes (Introduction). In this paper, we relate the proportions of different antibiotic types/classes among the overall volume of outpatient antibiotic prescription in different US states and English Clinical Commissioning Groups (CCGs) [37,39] to rates of mortality with sepsis in different age groups of US adults [38] and rates of *E. coli* and MSSA bacteremia in England [40]. Our results suggest, among other things, that prescribing of penicillins is associated with rates of *E. coli* and MSSA bacteremia in England, and rates of mortality with sepsis in older US adults, with the latter finding supporting our earlier results on the association between the use of penicillins and rates of septicemia hospitalization in older US adults [14]. We also note the high prevalence of resistance to penicillins in both the Gram-negative and Gram-positive infections [52–54]. Additionally, multivariable analyses of the US data suggest a positive association between the percent of individuals lacking health insurance and rates of mortality with sepsis in persons aged 50-64y, as well as the percent of individuals who are African-American and rates of mortality with sepsis in persons aged 65-84y, supporting the fact that rates of mortality with sepsis in African Americans are elevated [47]. While our results lend support for the replacement of penicillins by other antibiotics with the aim of reducing the rates of bacteremia/sepsis and associated mortality, more granular analyses, particularly individual-level studies relating prescribing of different antibiotics in the treatment of a given syndrome to subsequent outcomes are needed to inform guidelines for antibiotic, particularly penicillin replacement, as well as for reduction in antibiotic prescribing, as explained further in the next two paragraphs.

Our findings about the positive associations between the use of penicillins and mortality with sepsis in older US adults are in agreement with the fact that prevalence of resistance to penicillins, particularly in older adults, is high for a variety of infections with both Gram-negative and Gram-positive bacteria [52–54]. Negative associations between the proportion of cephalosporins among all antibiotic prescriptions and rates of sepsis mortality in older US adults (Table 3) may be related to the competition in prescribing with other antibiotic classes, particularly penicillins and fluoroquinolones for which prevalence of resistance in the key syndromes leading to sepsis is higher than prevalence of resistance to cephalosporins, e.g. [52,55]. We note that those negative associations do not reach statistical significance in the multivariable model (Table 4). Moreover, prevalence of cephalosporin resistance and the frequency of extended-spectrum beta-lactamase (ESBL) production, including in Gram-negative bacteria is growing [56], and replacement of other antibiotics by cephalosporins might potentially lead to negative effects in the long run. Finally, we found no associations between the proportion of macrolides among all antibiotic prescriptions in the US and rates of mortality with sepsis in adults, which agrees with our earlier findings regarding septicemia hospitalizations [14]. While macrolides are used relatively infrequently in the treatment of urinary tract and gastrointestinal infections, macrolides are commonly prescribed in the treatment of other sources of sepsis, particularly respiratory illness, both chronic [57] and acute, including pneumonia [58], with high prevalence of macrolide resistance in the corresponding infections [59].

In England, prevalence of resistance to trimethoprim in urinary tract infections (UTIs) is high [21], and trimethoprim use was found to be associated with UTI-related *E. coli* bacteremia [27]. Major reductions in trimethoprim prescribing in England took place in the recent years, particularly in 2017/2018 ([20]; Table 6 in this paper); moreover, prescribing of trimethoprim appears to have declined disproportionately in places in England with higher rates of *E. coli* and MSSA bacteremia (Results). All those changes might have played a role in the fact that growth in the rates of *E. coli* bacteremia in England has stalled in 2017/2018 after many years of robust increases (Table 5; [9,21,7,40]). Prevalence of co-amoxiclav resistance in *E. coli* bacteremia in England exceeds 40% [20,21], more than twice as high as the prevalence of co-amoxiclav resistance in *E.* coli-related urinary tract infections [21], suggesting that the use of co-amoxiclav and possibly of related penicillins is likely in the causal pathway for bacteremia outcomes. GP prescribing of co-amoxiclav was reduced significantly during the recent years ([20]; Table 6 in this paper), disproportionately in places with higher rates of bacteremia (Results). At the same time, prevalence of co-amoxiclav resistance in *E. coli* bacteremia and rates of the corresponding bacteremia outcomes are affected not only by GP prescribing of co-amoxiclav but also by other factors including the use of co-amoxiclav in secondary care, which is widespread [20,21], and possibly the use of related penicillins, with penicillin prescribing in secondary care increasing during the recent years [20]. We also note that amoxicillin use is associated with trimethoprim resistance [26], which in turn affects the rates of *E. coli* bacteremia [27], while use of different penicillins may also affect prevalence of resistance to piperacillin/tazobactam in *E. coli* bacteremia, which is sizeable [20]. Additionally, penicillins are widely prescribed in England, accounting for about half of all antibiotic prescriptions in primary care (e.g. Table 6), and penicillin use/resistance to penicillins is therefor expected to affect prevalence of infection/colonization with different bacterial pathogens that subsequently lead to bacteremia outcomes. While positive correlations between rates of bacteremia and rates of prescribing for penicillins other than amoxicillin/co-amoxiclav, but not for amoxicillin or co-amoxiclav were found in this paper, it is uncertain which penicillins (including co-amoxiclav and amoxicillin) have a greater relative impact on rates of bacteremia, with the results of the correlation analyses in this paper potentially affected by patterns of prescribing of different antibiotics related to different geographic locations, as well as demographic factors. Overall, our results support some replacement of penicillins in England by other antibiotics, presumably ones for which prevalence of antimicrobial resistance is lower, as well as reduction in penicillin prescribing with the aim of reducing bacteremia rates. Additionally, we have found that prescribing of metronidazole for skin infections (BNF 1310012K0) as a proportion of the overall GP antibiotic prescribing is correlated with the CCG-specific rates of both *E. coli* and MSSA bacteremia per unit of antibiotic prescribing for the four years in the data. This suggests the possibility that demographic/geographic differences result in differences in transmission of infections/colonization with bacterial pathogens such as *S. aureus* and *E. coli* (including transmission through skin infections), which in turn may affect the rates of severe bacterial infections, including bacteremia. Further work is needed to better understand those differences in transmission, including the feasibility of mitigation efforts aimed at preventing infections.

The epidemiological situation related to bacteremia/sepsis in England and the US brings about the question regarding the relative utility of antibiotic replacement vs. reduction in antibiotic prescribing for reducing the rates of bacteremia/sepsis and the associated mortality. A key mechanism relating antibiotic use to the rates of severe outcomes associated with bacterial infection is lack of clearance of resistant infections following antibiotic treatment, with some of those infections subsequently devolving into bacteremia/sepsis and lethal outcomes. While replacement of antibiotics (particularly penicillins) by those to which prevalence of resistance is lower should decrease the scale of this phenomenon, reduction in antibiotic prescribing without antibiotic replacement is not expected to bring down the rates of severe outcomes associated with bacterial infections, at least in the short term, as no treatment should generally be worse compared to antibiotic treatment with regard to sepsis-related outcomes. We note that several years of decreases in outpatient antibiotic prescribing in England before 2017/18 [23] did not seem to have an effect on the long term growth in the rates of both *E. coli* and MSSA bacteremia (Tables 6 and 5; [21,9,40]); at the same time, major replacement of trimethoprim by nitrofurantoin in 2017/18 (Table 6) following the issue of the corresponding guidelines for the treatment of UTIs ([21], p. 6) was accompanied by the stalling in the growth of the rates of *E. coli* bacteremia in England (Table 5 and [40]). Moreover, reduction in antibiotic prescribing has potential detrimental effects such as an increase in the volume of pneumonia hospitalization [24,25]. Reduction in antibiotic prescribing may contribute to decreases in the rates of severe outcomes associated with bacterial infections in the longer term through decreases in antibiotic resistance. While overall recommendations for reduction in antibiotic use are commonly issued by public health entities in different countries, e.g. [23], recommendations for replacement of certain antibiotics by certain others in the treatment of certain syndromes (like the recommendation for the replacement of trimethoprim by nitrofurantoin in England) are generally less common. Such recommendations related to penicillin use (e.g. in-hospital co-amoxiclav prescribing in England) should have a notable effect on the rates of bacteremia/sepsis and associated mortality, both in England and the US. Finally, we note that recently, US FDA has recommended the restriction of fluoroquinolone use for certain conditions (such as uncomplicated UTIs) due to potential adverse effects [60]. At the same time, no indications for antibiotics serving as replacement of fluoroquinolones were suggested in the FDA guidelines [60]. Such indications are needed to optimize the effect of those guidelines on the rates of severe outcomes associated with bacterial infections rather than possibly contribute to increases in the rates of such outcomes (e.g. increases in the rates of septicemia/sepsis through increases in the prescribing of penicillins).

Our paper has some limitations. Associations between proportions of different antibiotic types/classes (particularly penicillins) and rates of severe outcomes associated with bacterial infections may be affected not only by the relative contributions of the use of a unit of different antibiotics to the rates of those severe outcomes but also by patterns of antibiotic prescribing in different locations. We note that penicillins are prescribed for a wide variety of indications, both in England and the US, affecting prevalence of infection/colonization with different bacterial pathogens, and that there is high prevalence of resistance to penicillins in both the Gram-negative and Gram-positive bacteria in the US (e.g. [52–54]), and high prevalence of co-amoxiclav resistance in *E. coli* bacteremia in England [20], all of which supports our results about the relation between prescribing of penicillins and rates of severe outcomes associated with bacterial infections. The antibiotic-sepsis mortality associations that we found in the multivariable model estimate causal effects only if the model is well-specified and all confounders are accounted for in the analysis. To adjust for potential effects of unmeasured and residual confounding, we included random effects for the ten US Health and Human Services regions, which led to an improvement in the model fits. Moreover, results of the univariate and the multivariable analyses generally agree on the direction of the effect of different antibiotics on the rates of mortality with sepsis in the US (Tables 3 and 4). Further work involving more granular data, particularly individual-level analysis relating prescribing of different antibiotics in the treatment of a given syndrome to subsequent outcomes is needed to better ascertain the strength of the associations found in this paper. No hospital antibiotic prescribing data were available for this study, and in-hospital antibiotic prescribing is expected to have a significant impact on the rates of outcomes associated with severe bacterial infections. For example, given the very high prevalence of co-amoxiclav resistance in *E. coli* bacteremia and the high levels of co-amoxiclav prescribing in the secondary care setting in England [20,21], it is likely that co-amoxiclav prescribing in the secondary care has a significant effect on the incidence of *E. coli* bacteremia in England, both co-amoxiclav resistant and overall. Coding practices for sepsis on death certificates may vary by US state [61]. Additionally, data on outpatient antibiotic prescribing in the whole population [37] were related to age-specific rates of mortality with sepsis in the US [38], while in England, no age-specific prescribing or bacteremia data were available for this study. We expect that those sources of noise/incompatibility should generally reduce precision and bias the correlations towards null rather than create spurious associations.

We believe that despite those limitations, our results suggest that prescribing of certain antibiotics, particularly penicillins is associated with rates of *E. coli* and MSSA bacteremia in England and rates of mortality with sepsis in older US adults, with the latter result supporting our earlier findings about the association between the rates of prescribing of penicillins and rates of hospitalization with septicemia in older US adults [14]. Additionally, there is high prevalence of resistance to penicillins for a variety of bacterial infections both in the US and England [52–54,20,21]. While these findings support the potential utility of replacement of penicillins by other antibiotics with the goal of reducing the rates of bacteremia/sepsis and associated mortality, further studies, including individual-level analyses are needed to better understand the effect of replacement of certain antibiotics, particularly penicillins by other antibiotics in the treatment of different syndromes, well as the effect of reduction in antibiotic prescribing in the treatment of certain conditions on the rates of severe outcomes associated with bacterial infections.

## Acknowledgment

We thank Koen Pouwels for helpful discussions.

